# DanQ: a hybrid convolutional and recurrent deep neural network for quantifying the function of DNA sequences

**DOI:** 10.1101/032821

**Authors:** Daniel Quang, Xiaohui Xie

## Abstract

Modeling the properties and functions of DNA sequences is an important, but challenging task in the broad field of genomics. This task is particularly difficult for noncoding DNA, the vast majority of which is still poorly understood in terms of function. A powerful predictive model for the function of noncoding DNA can have enormous benefit for both basic science and translational research because over 98% of the human genome is noncoding and 93% of disease-associated variants lie in these regions. To address this need, we propose DanQ, a novel hybrid convolutional and bi-directional long short-term memory recurrent neural network framework for predicting noncoding function *de novo* from sequence. In the DanQ model, the convolution layer captures regulatory motifs, while the recurrent layer captures long-term dependencies between the motifs in order to learn a regulatory “grammar” to improve predictions. DanQ improves considerably upon other models across several metrics. For some regulatory markers, DanQ can achieve over a 50% relative improvement in the area under the precision-recall curve metric compared to related models.

Availability and implementation

All source code is available at the github repository http://github.com/uci-cbcl/DanQ.

## Introduction

The recent deluge of high throughput genomic sequencing data has prompted the development of novel bioinformatics algorithms that can integrate large, feature-rich datasets. Deep learning algorithms are attractive solutions for such problems because they are scalable with large datasets and are effective in identifying complex patterns from feature-rich datasets (LeCun et al., 2015). They are able to do so because deep learning algorithms utilize heuristics and specialized hardware to efficiently train deep neural networks (DNNs) that learn high levels of abstractions from multiple layers of non-linear transformations. DNNs have already been adapted for problems such as motif discovery (Alipanahi et al., 2015) and predicting the deleteriousness of genetic variants (Quang et al., 2015).

There has been a growing interest to predict function directly from sequence, instead of from curated datasets such as gene models and multiple species alignment. Much of this interest is attributed to the fact that over 98% of the human genome is noncoding, the function of which is not very well-defined. A model that can predict function directly from sequence may reveal novel insights about these noncoding elements. Over 1,200 genome-wide association studies have identified nearly 6,500 disease- or trait-predisposing SNPs, 93% of which are located in noncoding regions (Hindorff et al., 2014), highlighting the importance of such a predictive model. Convolutional neural networks (CNNs) are variants of DNNs that are appropriate for this task. CNNs use a weight-sharing strategy to capture local patterns in data such as sequences. This weight-sharing strategy is especially useful for studying DNA because the convolution filters can capture sequence motifs, which are short, recurring patterns in DNA that are presumed to have a biological function. DeepSEA is a recently developed algorithm that utilizes a CNN for predicting DNA function (Zhou and Troyanskaya, 2015). The CNN is trained in a joint multi-task fashion to simultaneously learn to predict large-scale chromatin-profiling data, including transcription factor (TF) binding, DNase I sensitivity and histone-mark profiles across multiple cell types, allowing the CNN to learn tissue-specific functions. It significantly outperforms gkm-SVM (Ghandi et al., 2014), a related algorithm that can also predicts the regulatory function of DNA sequences, but uses a support vector machine instead of a CNN for predictions. To predict the effect of regulatory variation, both gkm-SVM (Lee et al., 2015) and DeepSEA use a similar strategy of predicting the function of both the reference and allele sequences and processing the score differences.

Another variation of DNNs is the recurrent neural network (RNN). Unlike a CNN, connections between units of a RNN form a directed cycle. This creates an internal state of the network that allows it to exhibit dynamic spatial behavior. A bi-directional long short-term memory network (BLSTM) is a variant of the RNN that combines the outputs of two RNNs, one processing the sequence from left to right, the other one from right to left. Instead of regular hidden units, the two RNNs contain LSTM blocks, which are “smart” network units that can remember a value for an arbitrary length of time. BLSTMs can capture long-term dependencies and have been effective for other machine learning applications such as phoneme classification (Graves and Schmidhuber, 2005), speech recognition (Graves et al., 2013), machine translation (Sundermeye et al., 2014), human action recognition (Zhu et al., 2015), and sentiment analysis (Li et al., 2015). Although BLSTMs are effective for studying sequential data, they have not been applied for DNA sequences.

Hence, we propose DanQ, a hybrid framework that combines CNNs and BLSTMs (Fig. 1). The first layers of the DanQ are designed to scan sequences for motif sites through convolution filtering. Whereas the convolution step of the DeepSEA model contains three convolution layers and two max pooling layers in alternating order to learn motifs, the convolution step of the DanQ model is much simpler and contains one convolution layer and one max pooling layer to learn motifs. The max pooling layer is followed by a BLSTM layer. Our rationale for including a recurrent layer after the max pooling layer is that motifs can follow a regulatory “grammar” governed by physical constraints that dictate the *in vivo* spatial arrangements and frequencies of combinations of motifs, a feature associated with tissue-specific functional elements such as enhancers (Quang and Xie, 2014; Quang et al., 2015b). Following the BLSTM layer, the last two layers of the DanQ model are a dense layer of rectified linear units and a multi-task sigmoid output, similar to the DeepSEA model.

**Figure 1.**
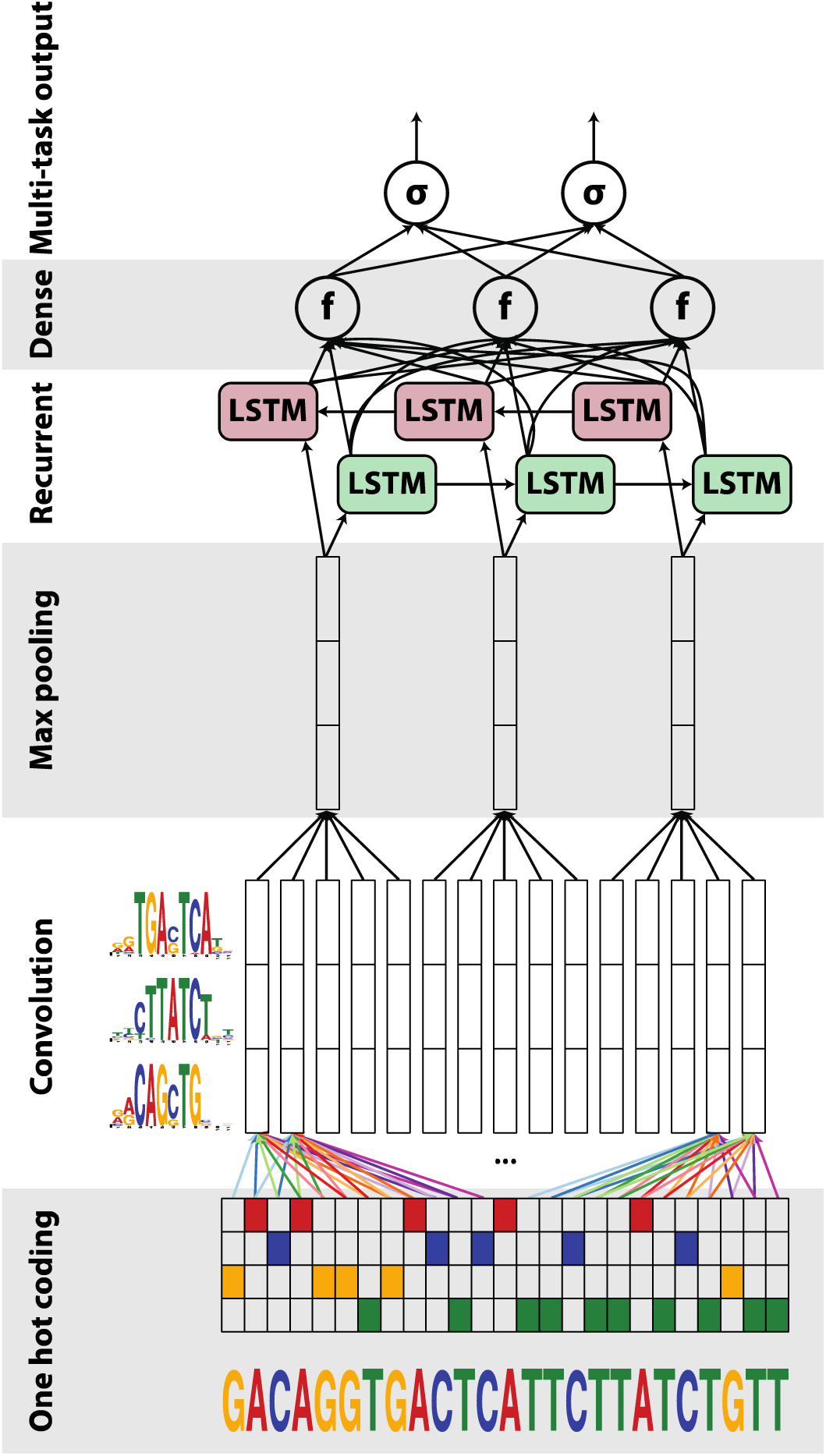
A graphical illustration of the DanQ model: An input sequence is first one hot encoded into a 4-row bit matrix. A convolution layer with rectifier activation acts as a motif scanner across the input matrix to produce an output matrix with a row for each convolution kernel and a column for each position in the input (minus the width of the kernel). Max pooling reduces the size of the output matrix along the spatial axis, preserving the number of channels. The subsequent BLSTM layer considers the orientations and spatial distances between the motifs. BLSTM outputs are flattened into a layer as inputs to a fully connected layer of rectified linear units. The final layer performs a sigmoid nonlinear transformation to a vector that serves as probability predictions of the epigenetic marks to be compared via a loss function to the true target vector.

DanQ surpasses other methods for predicting the properties and function of DNA sequences across several metrics. In addition, we show that the convolution kernels learned by the model can be converted to motifs, many of which significantly match known motifs. We expect DanQ to provide novel insights into noncoding genomic regions and contribute to understanding the potential functions of complex disease- or trait-associated genetic variants.

## Methods

### Features and data

The DanQ framework uses the same features and data as the DeepSEA framework. Briefly, the human GRCh37 reference genome was segmented into non-overlapping 200-bp bins. Targets were computed by intersecting 919 ChIP-seq and DNase-seq peak sets from uniformly processed ENCODE and Roadmap Epigenomics data releases, yielding a length 919 binary target vector for each sample. Each sample input consists of a 1,000-bp sequence centered on a 200-bp bin that overlaps at least one TF binding ChIP-seq peak, and is paired with the respective target vector. Based on this information, we expected that each target vector would contain at least one positive value; however, we found that about 10% of all target vectors were all negatives. Each 1,000-bp DNA sequence is one-hot encoded into a 1,000 × 4 binary matrix, with columns corresponding to A, G, C and T. Training, validation, and testing sets were downloaded from http://deepsea.princeton.edu/media/code/deepsea_train_bundle.v0.9.tar.gz. Samples were stratified by chromosomes into strictly nonoverlapping training, validation and testing sets. The validation set was not used for training or testing. Reverse complements are also included, effectively doubling the size of each dataset.

For evaluating performance on the test set, the predicted probability for each sequence was computed as the average of the probability predictions for the forward and reverse complement sequence pairs, similar to DeepSEA’s evaluation experiments.

### DanQ model and training

For detailed specifications of the architectures and hyperparameters used in this study, see Supplementary Note. Dropout (Srivastava et al., 2014) is included to randomly set a proportion of neuron activations from the max pooling and BLSTM layers to a value of 0 in each training step to regularize the DanQ models.

All weights are initialized by randomly drawing from U(−0.05,0.05) and all biases are initially set to 0. In addition to random initialization, an alternative strategy is to initialize kernels from known motifs: a random subsection of a kernel is set equal to the values of the position frequency matrix minus 0.25, and its corresponding bias is randomly drawn from U(−1.0,0.0). We tried both in our implementation.

Neural network models are trained using the RMSprop algorithm (Tielemanand Hinton 2012) with a minibatch size of 100 to minimize the average multi-task binary cross entropy loss function on the training set. Validation loss is evaluated at the end of each training epoch to monitor convergence. The first model we trained contains 320 convolution kernels with random initial weights, referred to as DanQ, took 60 epochs to fully train, and each epoch of training takes approximately 6 hours. The second model we trained, which we designated as DanQ-JASPAR because half of the kernels are initialized with motifs from the JASPAR database (Mathelier et al., 2015), contains 1024 convolution kernels, took 30 epochs to fully train, and each epoch of training takes approximately 12 hours.

Our implementation utilizes the Keras 0.2.0 library (https://github.com/fchollet/keras) with the Theano 0.7.1 (Bastien et al., 2012; Bergstra et al., 2010) backend. An NVIDIA Titan Z GPU was used for training the model.

### Logistic regression

For benchmark purposes, we also trained a logistic regression (LR) baseline model. Unlike the DanQ and DeepSEA models, the LR model does not process raw sequences as inputs. Instead, the LR model uses zero-mean and unit variance normalized counts of k-mers of lengths 1–5bp as features. The LR model was regularized with a small L2 weight regularization of 1e-6. Similar to the training of the DanQ models, the LR model was trained using the RMSprop algorithm with a minibatch size of 100 to minimize the average multi-task binary cross entropy loss function on the training set. Validation loss is evaluated at the end of each training epoch to monitor convergence. We note that this method of training is equivalent to training 919 individual single-task LR models.

## Results

We first train a DanQ model containing 320 convolution kernels for 60 epochs, evaluating the average multi-task cross entropy loss on the validation set at the end of each epoch to monitor the progress of training. To regularize the model, we also include dropout to randomly set a proportion of neuron activations from the max pooling and BLSTM layers to a value of 0 in each training step. For detailed specifications of the hyperparameters and model architecture, see Supplementary Note.

For benchmarking purposes, we compare a fully trained DanQ model to a LR baseline model and the published DeepSEA model. To compare performance among models, we first calculated the area under the receiver operating characteristics curve (ROC AUC) for each of the 919 binary targets on the test set (Fig. 2). In terms of the ROC AUC score, DanQ outperforms the DeepSEA model for two of the targets as shown in the examples at the top of Fig. 2, although this performance difference is relatively small. This pattern extends to the remaining targets as DanQ outperforms DeepSEA for 94.1% of the targets, although the difference is again comparatively small with an absolute improvement of around 1–4% for most targets. Despite the simplicity of the LR models, the ROC AUC statistics suggests that LR is an effective predictor, with ROC AUC scores typically over 70%. Given the sparsity of positive binary targets (~2%), the ROC AUC statistic is highly inflated by the class imbalance, a fact overlooked in the original DeepSEA paper.

**Figure 2.**
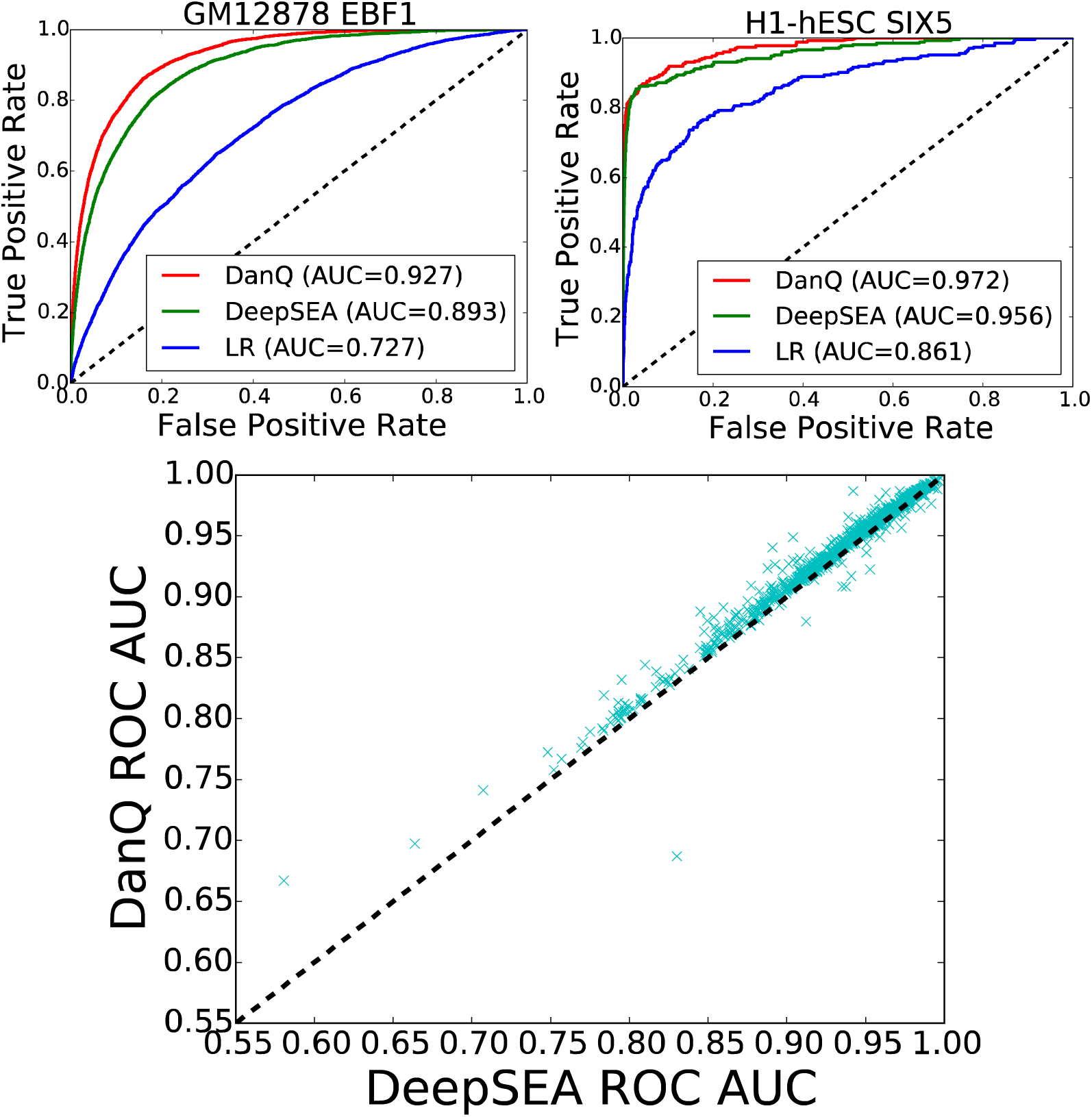
(top) ROC curves for the GM12S7S EBF1 and H1-hESC SIX5 targets comparing the performance of the three models. (bottom) A direct comparison of ROC AUC between DanQ and DeepSEA for each of the 919 targets, showing that DanQ outperforms DeepSEA for 94.1% of the targets in terms of ROC AUC.

A better metric to measure the performance is the area under precision-recall curve (PR AUC) (Fig. 3). Neither the precision nor recall take into account the number of true negatives, thus the PR AUC metric is less prone to inflation by the class imbalance than the ROC AUC metric is. As expected, we found the PR AUC metric to be more balanced, as demonstrated by how the LR models now achieve a PR AUC below 5% for the two examples at the top of Fig. 3, far below the performance of the other two models. Moreover, the performance gap between DanQ between DeepSEA is much more pronounced under the PR AUC statistic than under the ROC AUC statistic. For the two examples shown, the absolute improvement is over 10% and the relative improvement is over 50% under the PR AUC metric, and 97.6% of all DanQ PR AUC scores surpass DeepSEA PR AUC scores. These results show that adding recurrent connections significantly increases the modeling power of DanQ.

**Figure 3.**
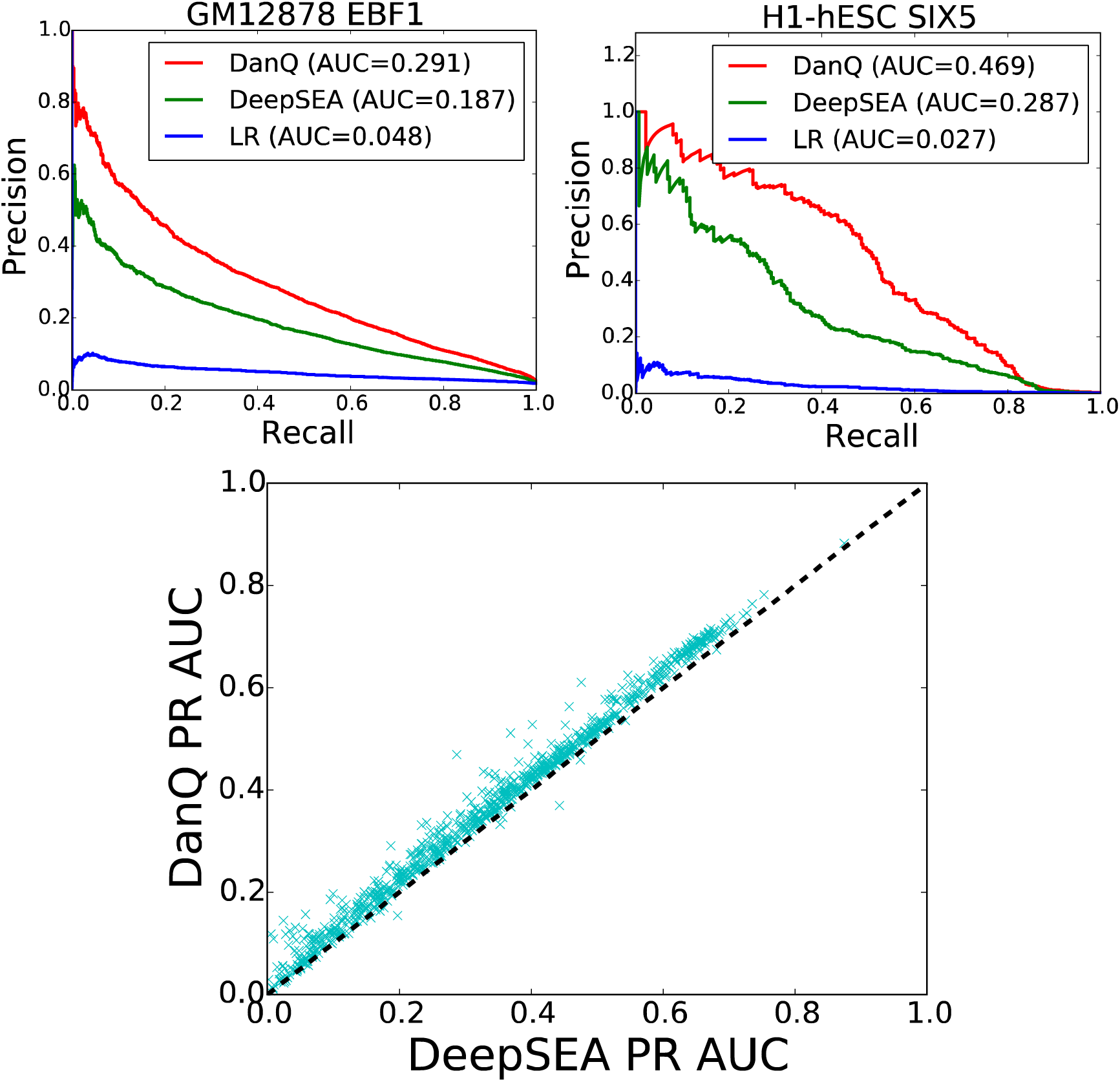
(top) PR curves for the GM12S7S EBF1 and H1-hESC SIX5 targets comparing the performance of the three models. (bottom) A direct comparison of PR AUC between DanQ and DeepSEA for each of 919 targets, showing that DanQ outperforms DeepSEA for 97.6% of the targets in terms of PR AUC.

Using a similar approach described in the DeepBind method (Alipanahi et al., 2015), we converted the kernels from the convolution layer of the DanQ models to position frequency matrices, or motifs. Then, we aligned these motifs to known motifs using the TOMTOM algorithm (Gupta et al., 2007). Of the 320 motifs learned by the DanQ model, 166 significantly match known motifs (*E*<0.01) (Fig. 4A, Fig S1, Supplemental File). Next, we aligned and clustered the 320 motifs together into 118 clusters using the RSAT matrix clustering tool (Medina-Rivera et al., 2015), and confirmed that the model learned a large variety of informative motifs (Fig. 4B and 4C).

**Figure 4.**
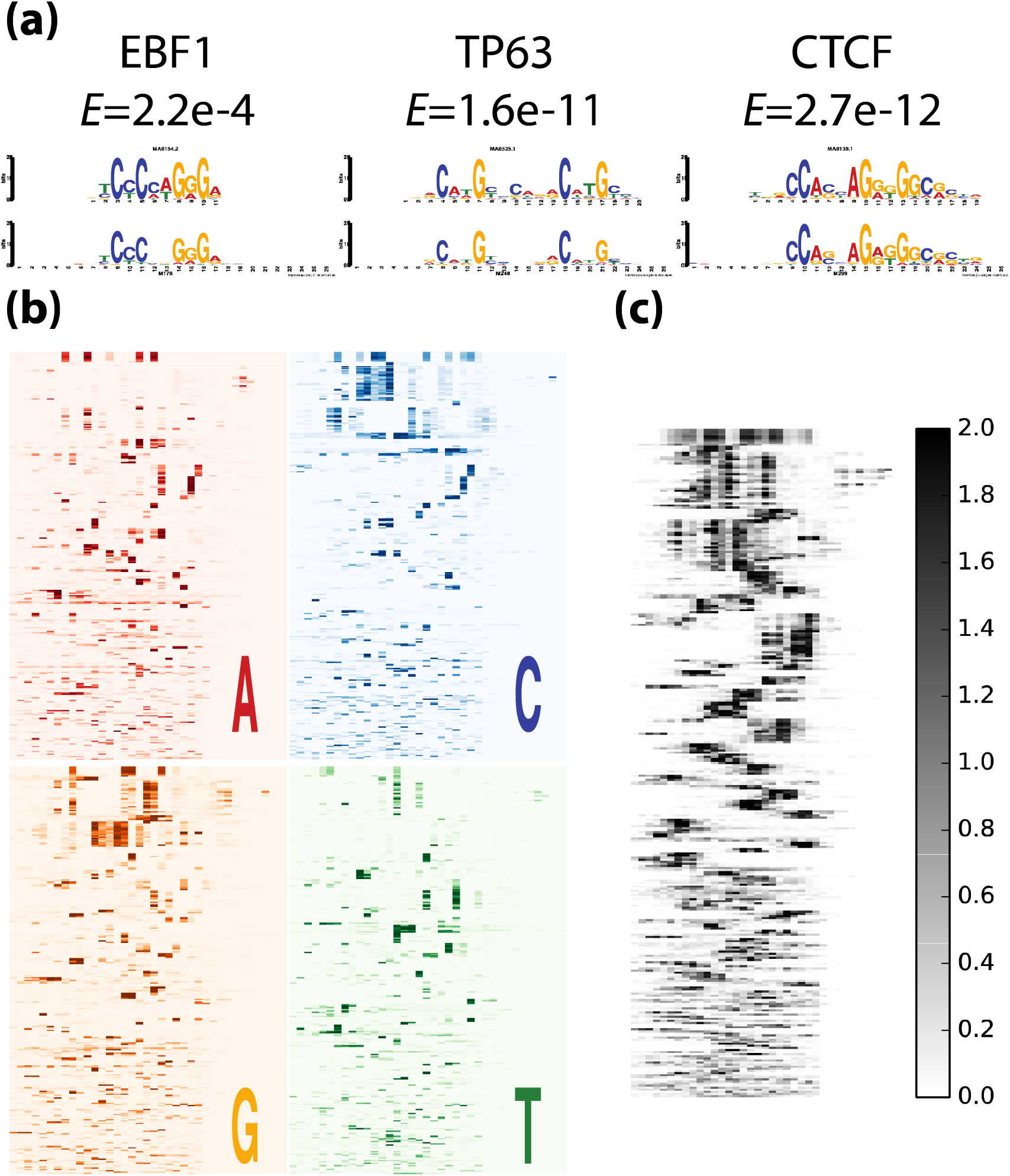
(a) Three convolution kernels (top) visualized and aligned with EBF1, TP63, and CTCF motif logos (bottom) from JASPAR using TOMTOM. Significance values of the match are displayed below motif names. (b) All 320 convolution kernels are converted to sequence logos and aligned with RSAT. The heatmaps are colored according to the information content of the respective nucleotide at each position. (c) Same as (b), except that the heatmap is colored by the sum of the information content of each letter.

Given the large scope of the data, we conjectured that our current model did not exhaust the entire space of useful motif features despite the large variety of motifs learned. Moreover, weight initialization is known to play crucial role for the performance neural networks (Sutskever et al., 2013) and we hypothesized that a better initialization strategy can further improve the performance of our neural network. Therefore, we trained a larger model containing 1024 convolutional kernels of which about half are initialized with known motifs from JASPAR (Mathelier et al., 2015) and found this alternative way of initialization can further improve the performance of DanQ (Table S1, Fig. S2). We are actively refining the architecture and initialization, and we will provide regular updates on the best model. We will also provide motifs from these models in MEME minimal format, a flexible format compatible with most motif-related programs, as a resource to the community.

## Discussion

In conclusion, DanQ is a powerful method for predicting the function of DNA directly from sequence alone, making it a valuable asset for studying the function of noncoding DNA. Its hybrid architecture allows it to simultaneously learn motifs and a complex regulatory “grammar” between the motifs. The additional modeling capacity afforded by the recurrent connections allows DanQ to significantly outperform DeepSEA, a pure CNN model that lacks recurrent modeling. This performance gap is demonstrated across several metrics, including a direct comparison of AUC statistics between the two models. We argue that the PR AUC statistic is a much more balanced metric than the ROC AUC statistic to assess performance in this case due to the massive class imbalance. In fact, the performance gap can be quite drastic under the PR AUC statistic, reaching well over a 50% relative improvement for some epigenetic marks. Nevertheless, despite the significant improvement in performance, there is still much room for improvement because most of the PR AUC scores are below 70% for either model.

There are several avenues of future interest to explore. First, we are actively exploring how to extend the model to process genetic variants in order to predict their functional consequences. Second, the model can be made fully recurrent so it can process sequences of arbitrary length, such as whole chromosome sequences, to generate sequential outputs. In contrast, our current setup can only processes sequences of constant length with static outputs. A fully recurrent architecture may also benefit our effort to study variants since it would allow us to explore the long-range consequences of genetic variants, as well as the cumulative effects of SNPs that are in linkage disequilibrium with each other. Finally, we are committed to updating and improving the DanQ model. This involves improving the model architecture and incorporating new ChIP-seq and DNase-seq datasets from more cell types as they become available. We are also interested in incorporating other types of data, such as methylation, nucleosome positioning, and even possibly transcription. To the best of our knowledge, this is the first application of a hybrid convolution and recurrent network architecture for the purpose of predicting function *de novo* from DNA sequences. We expect this hybrid architecture will be continually explored for the purpose of studying biological sequences.

## Funding

National Institute of Biomedical Imaging and Bioengineering, National Research Service Award (EB009418) from the University of California, Irvine, Center for Complex Biological Systems. This material is based upon work supported by the National Science Foundation Graduate Research Fellowship under Grant No. (DGE-1321846). Any opinion, findings, and conclusions or recommendations expressed in this material are those of the authors(s) and do not necessarily reflect the views of the National Science Foundation.

## Acknowledgements

We gratefully acknowledge Jian Zhou for helping us understand DeepSEA and providing the datasets.Conflict of Interest: none declared.

## Supplementary note

Detailed specifications of the architectures and hyperparameters of the DanQ models used in this study. Numbers to the right of the forward slash indicate values unique to the DanQ-JASPAR model, a larger model in which about half of the convolution kernels are initialized with motifs from the JASPAR database.

Model Architecture:
1. Convolution layer (320/1024 kernels. Window size: 26/30. Step size: 1.)
2. Pooling layer (Window size: 13/15. Step size: 13/15.)
3. Bi-directional long short term memory layer (320/512 forward and 320/512 backward LSTM neurons)
4. Fully connected layer (925 neurons)
5. Sigmoid output layer

Regularization Parameters:

Dropout proportion (proportion of outputs randomly set to 0):

Layer 2: 20% Layer 3: 50% All other layers: 0%

**Table S1.** Accuracy and cross-entropy loss on training, validation, and testing sets for each of the models. The DanQ model initialized with JASPAR motifs performed the best across all metrics, as indicated in bold. Note that due to the huge class imbalance, all models achieved high accuracies.

|  | Training |  | Validation |  | Testing |  |
| --- | --- | --- | --- | --- | --- | --- |
|  | Loss | Accuracy | Loss | Accuracy | Loss | Accuracy |
| Predict all 0's | N/A | 97.84% | N/A | 98.05% | N/A | 97.94% |
| LR | 0.0798 | 97.90% | 0.0743 | 98.09% | 0.0771 | 97.99% |
| DeepSEA | 0.0551 | 98.20% | 0.0509 | 98.36% | 0.0554 | 98.21% |
| DanQ | 0.0539 | 98.23% | 0.0491 | 98.39% | 0.0538 | 98.24% |
| **DanQ-JASPAR** | **0.0521** | **98.26%** | **0.0482** | **98.40%** | **0.0533** | **98.24%** |

**Figure S1.**
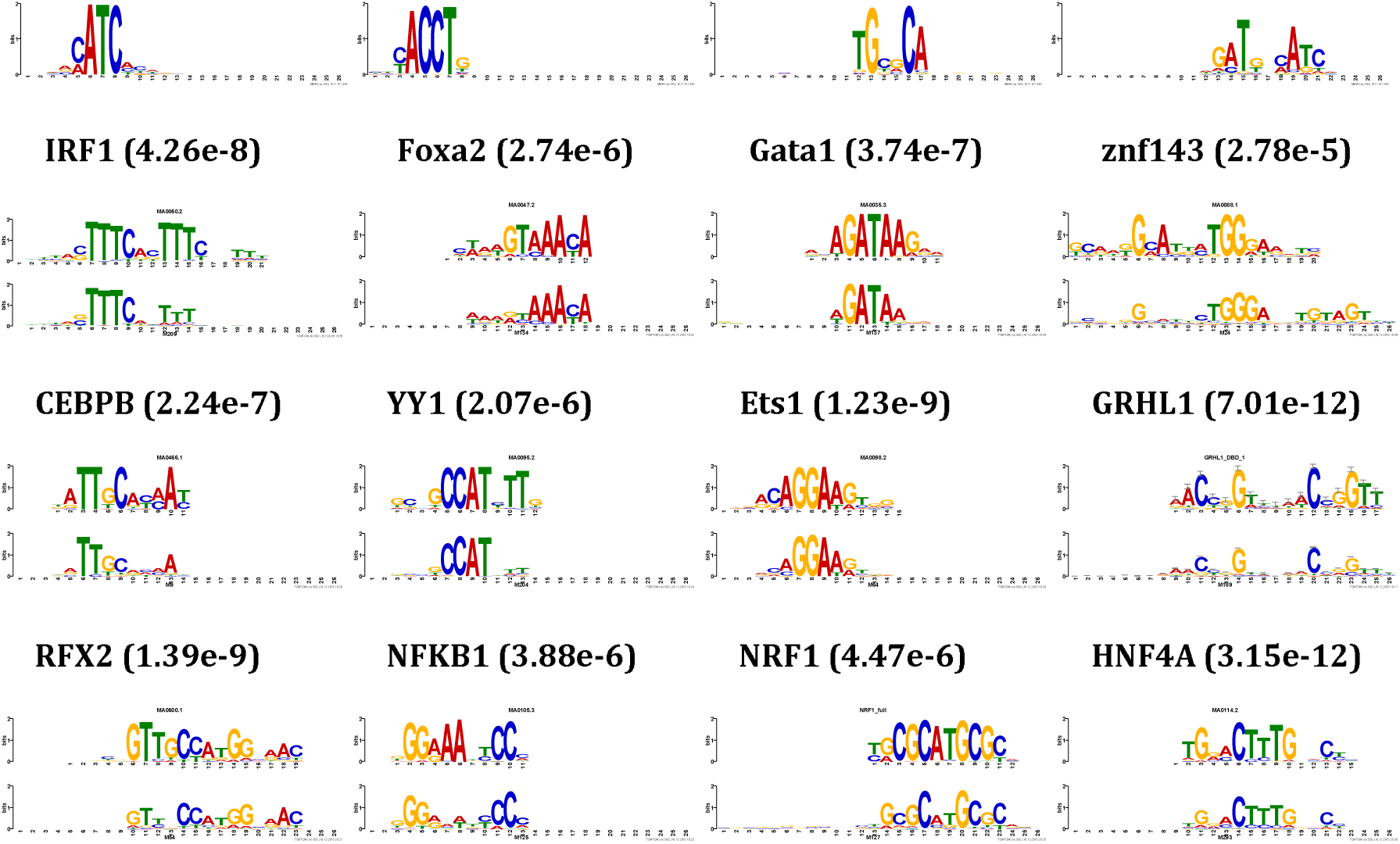
Examples of convolution kernels from the DanQ model converted to motif logos. The top four motifs did not significantly match (E < 0.01) any vertebrate-related motifs according to TOMTOM. Each of the remaining twelve panels shows the logo of the motif discovered by DanQ (lower logo) aligned with the best matching motif in the databases (upper logo), along with the name of the best matching transcription factor and significance value of the match.

**Figure S2.**
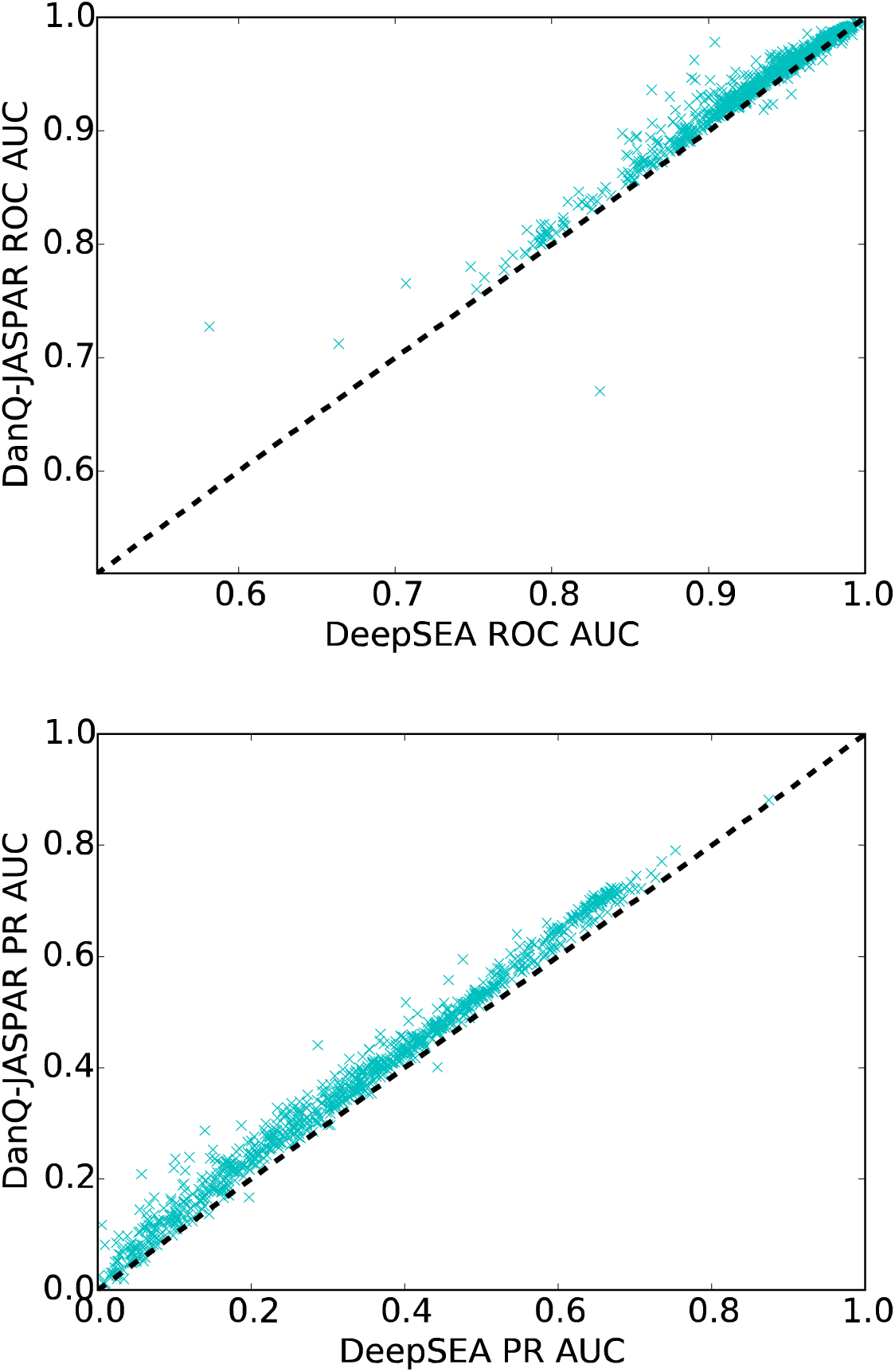
A comparison of ROC and PR AUC statistics for each of the 919 binary targets between the DanQ-JASPAR and DeepSEA models. 96.6% and 98.6% of ROC and PR AUC scores are greater for the DanQ-JASPAR model than they are for the DeepSEA model, respectively.

